# Normal Olfactory Functional Connectivity Despite Life-Long Absence of Olfactory Experiences

**DOI:** 10.1101/2020.05.21.106161

**Authors:** Moa G. Peter, Peter Fransson, Gustav Mårtensson, Elbrich M. Postma, Love Engström Nordin, Eric Westman, Sanne Boesveldt, Johan N. Lundström

## Abstract

Congenital blindness is associated with atypical morphology, and functional connectivity within and from, visual cortical regions; changes that are hypothesized to originate from a life-long absence of visual input and could be regarded as a general (re)organization principle of sensory cortices. Challenging this is the fact that individuals with congenital anosmia (life-long olfactory sensory loss) display little to no morphological changes in primary olfactory cortex. To determine whether olfactory input from birth is essential to establish and maintain normal functional connectivity in olfactory processing regions, akin to the visual system, we assessed differences in functional connectivity within olfactory cortex between individuals with congenital anosmia (n=33) and matched controls (n=34). Specifically, we assessed differences in connectivity between core olfactory processing regions as well as differences in regional homogeneity and homotopic connectivity within primary olfactory cortex. In contrast to congenital blindness, none of the analyses indicated atypical connectivity in individuals with congenital anosmia. In fact, post-hoc Bayesian analysis provided support for an absence of group differences. These results suggest that a lifelong absence of olfactory experience has limited impact on the functional connectivity in olfactory cortex, a finding that indicates a clear difference between sensory modalities in how sensory cortical regions develop.

## INTRODUCTION

Absence of input from a sensory modality has been linked to notable alterations in the human brain (Bavelier and Neville 2002; Merabet and Pascual-Leone 2010; Frasnelli et al. 2011). These alterations consist of, e.g., morphological changes within cortical areas normally devoted to the processing of the lost sense (Park et al. 2009; Emmorey et al. 2003), altered processing of input from the remaining senses (Sadato et al. 2005; Collignon et al. 2013), and are often linked to behavioral changes (Voss and Zatorre 2012; Gougoux et al. 2005). However, in comparison to auditory and visual sensory deprivation, what effect congenital anosmia (a lifelong lack of olfactory input) has on cortical regions vital for olfactory processing has been scarcely studied and is poorly understood. That olfactory loss is poorly studied is not surprising given that many think, erroneously, that the human sense of smell is a residual sense; however, humans have a better sense of smell than most animals (McGann 2017) and olfactory information is of importance for our wellbeing (Croy et al. 2014). Congenital anosmia has been linked to morphological reorganization within the orbitofrontal cortex (Frasnelli et al. 2013; Karstensen et al. 2018; Peter et al. 2019) including areas commonly referred to as secondary olfactory cortex (Lundström et al. 2011), but, surprisingly, it has been suggested that no morphological alterations occur within primary olfactory cortex despite a life-long absence of olfactory input (Peter et al. 2019). As of today, it is not known whether these morphological deviations (and suggested lack thereof) are reflected in functional neuroimaging data. By assessing potential functional anomalies in the olfactory network in individuals with congenital anosmia, we can gain a better understanding of whether olfactory sensory experience is critical for normal development of the olfactory brain.

Assessing functional activity within a sensory neural network without presenting the associated sensory stimuli is problematic. However, fluctuations in the blood-oxygen-level dependent (BOLD) signal at rest, measured by resting-state functional magnetic resonance imaging (fMRI), have been linked to intrinsic functional networks in the adult brain (Damoiseaux et al. 2006; Biswal et al. 1995; Fransson 2005). Hence, resting-state fMRI allows us to assess how the lack of sensory input during development influence connectivity in sensory processing regions without requiring sensory input. Both visual and auditory congenital sensory deprivation has repeatedly been linked to altered resting-state functional connectivity within early sensory processing regions of the respective deprived sense as well as altered inter-regional connectivity (Wang et al. 2014; Ding et al. 2016). Specifically, the visual cortex of blind individuals shows a higher regional homogeneity of the BOLD signal at rest (A. Jiang et al. 2015; Liu et al. 2011), i.e., a higher similarity of BOLD signal in neighboring voxels, as well as a lower homotopic connectivity (Watkins et al. 2012; Hou et al. 2017), in other words a lower interhemispheric functional connectivity, as compared to controls. Because regional homogeneity typically decreases and homotopic connectivity typically increases in sensory processing areas during normal development (Zuo et al. 2010; Anderson et al. 2014), these anomalies in blind individuals suggest that sensory input might drive the development and preservation of functional connectivity.

To determine whether olfactory experience is critical to develop normal functional connectivity within the olfactory system, we assess what effects a lifelong absence of olfactory input has on the functional connectivity within and between core olfactory regions by comparing individuals with isolated congenital anosmia (ICA; individuals with congenital anosmia that cannot be ascribed to a specific event or condition) to age, sex, and education matched individuals with established functional olfactory abilities. First, we will determine potential ICA-dependent differences in functional connectivity within the olfactory network by comparing resting-state functional connectivity between regions important for olfactory processing. Here, we hypothesize that the life-long absence of olfactory input in individuals with ICA is associated with a lower functional connectivity between said regions. Second, we will determine whether individuals with ICA demonstrate functional alterations within primary olfactory cortex by assessing regional homogeneity and homotopic connectivity. Based on previously published effects of visual sensory loss, we hypothesize that the primary olfactory cortex in individuals with ICA shows higher regional homogeneity and lower homotopic connectivity.

## METHODS

### Participants

A total of 68 individuals participated: 34 individuals with ICA and 34 controls matched in terms of age, sex, and educational level (Table 1). One individual with ICA was excluded in the post-scanning image quality control phase due to the detection of a morphological anomaly.

**Table 1.**
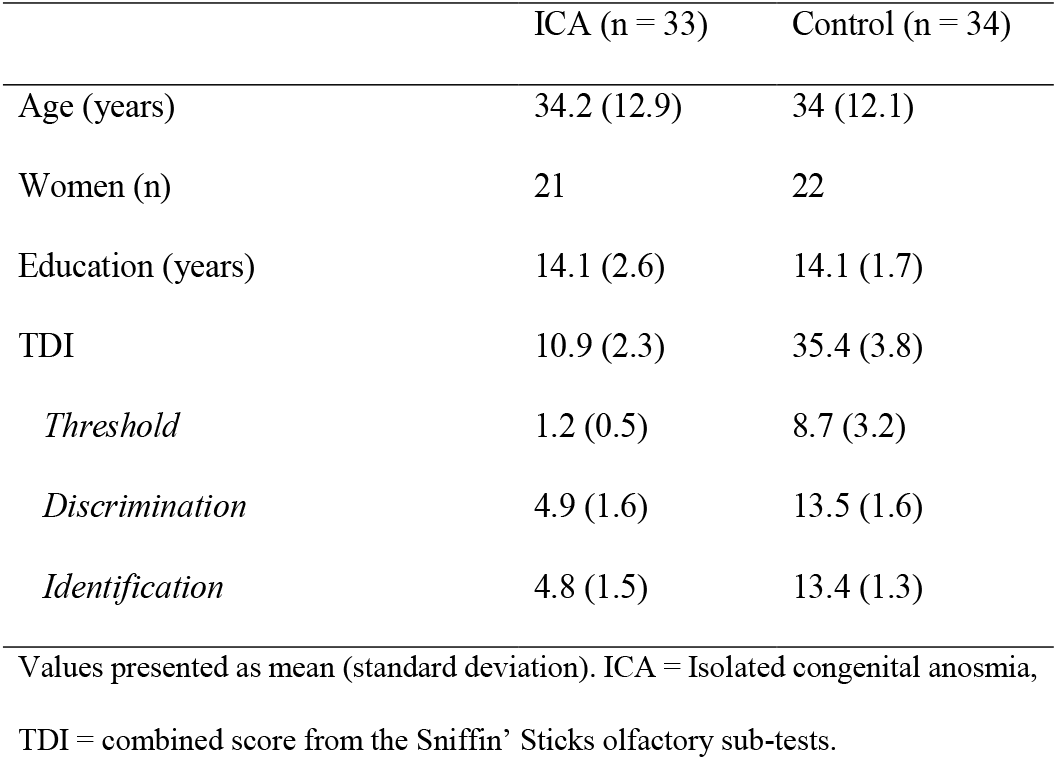
Descriptive statistics per experimental group.

|  | ICA (n = 33) | Control (n = 34) |
| --- | --- | --- |
| Age (years) | 34.2 (12.9) | 34 (12.1) |
| Women (n) | 21 | 22 |
| Education (years) | 14.1 (2.6) | 14.1 (1.7) |
| TDI | 10.9 (2.3) | 35.4 (3.8) |
| *Threshold* | 1.2 (0.5) | 8.7 (3.2) |
| *Discrimination* | 4.9 (1.6) | 13.5 (1.6) |
| *Identification* | 4.8 (1.5) | 13.4 (1.3) |

Values presented as mean (standard deviation). ICA = Isolated congenital anosmia,
TDI = combined score from the Sniffin' Sticks olfactory sub-tests.

Diagnosing ICA is a non-trivial problem due to the difficulty of dissociating a congenital cause from a very early onset of anosmia. We took a number of precautions to decrease the risk of incorrectly including an individual in the ICA group. All participants labeled as ICA reported a lifelong absence of olfactory perception and, based on questionnaires, absence of potential non-congenital causes of their anosmia (e.g. head trauma, disease, exposure to toxins), or other congenital disorders (e.g. Kallmann syndrome). Additionally, 24 of the individuals with ICA had previously sought medical care for their inability to smell odors and received the clinical diagnose anosmia. Finally, all participants were tested to establish normal sense of smell (controls) or functional anosmia (ICA). Due to the rareness of ICA, participants were recruited and tested at two different sites: 46 individuals (23 ICA) in Stockholm, Sweden, and 22 individuals (11 ICA) in Wageningen, the Netherlands; a matched control was always scanned at the same site as an individual with ICA. The study was approved by the ethical review boards in both Sweden and in the Netherlands and all participants provided written informed consent prior to study participation.

### Olfactory screening

Participants’ olfactory ability was assessed using the full Sniffin’ Sticks olfactory test battery (Burghart, Wedel, Germany). The test consists of three subtests assessing odor detection threshold (T), odor quality discrimination (D), and cued odor quality identification (I), each rendering a maximum score of 16. The combined score of the subtests (TDI) was used to assess individual olfactory ability compared to normative data from over 3000 individuals (Hummel et al. 2007), corresponding well with a recent publication including 9000 individuals (Oleszkiewicz et al. 2019). Individuals with ICA had a mean TDI score of 10.9 (SD = 2.3, range: 7-15; all below the limit for functional anosmia) and control individuals had a mean TDI score of 35.4 (SD = 3.8, range: 28.5-42.5; all above the 10th percentile within their respective age and sex group; Table 1).

### Image acquisition

Imaging data was acquired on two 3T Siemens Magnetom MR scanners (Siemens Healthcare, Erlangen, Germany): a Prisma scanner using a 20-channel head coil (Sweden) and a Verio scanner using a 32-channel head coil (the Netherlands). Both sites used identical scanning sequence protocols and participant instructions.

A 9 minutes long resting-state scan was acquired for each participant using an echo-planar imaging sequence (TR = 2000 ms, TE = 22 ms, flip angle = 70°, voxel size = 3·3·3 mm3, 41 slices, FOV = 228 mm, interleaved acquisition). To reduce the risk of participants falling asleep, participants were instructed to keep their eyes open and look at a fixation cross. A structural T1-weighted image was acquired for each individual using an MP-RAGE sequence (TR = 1900 ms, TI = 900 ms, TE = 2.52 ms, flip angle = 9°, voxel size = 1·1·1 mm3, 176 slices, FOV = 256 mm). Results from structural image analysis are reported elsewhere (Peter et al. 2019).

### Image analyses

#### Preprocessing

The data was preprocessed using SPM12 software (Wellcome Trust Centre for Neuroimaging, UCL; https://www.fil.ion.ucl.ac.uk/spm/) running in MATLAB 2019b (The MathWorks, Inc., Natick, Massachusetts, USA). The preprocessing pipeline included slice timing correction, image realigning using a 6 parameter rigid body transformation model, coregistering the structural image to the mean functional image using affine transformation, and normalizing both structural and functional images to MNI-space using functions based on the “unified segmentation” approach (Ashburner and Friston 2005). The structural image was segmented, bias corrected, and spatially normalized, and the deformation field for normalizing the structural image was thereafter used to normalize the functional images.

#### Denoising

Motion artifacts are particularly problematic in functional connectivity analysis of resting state fMRI. To reduce the likelihood of bias of the results from head-motion, Power’s frame-wise displacement measure (Power et al. 2012) was used to visually inspect motion, statistically compare motion between the two groups, and decrease the influence of motion-related noise by data scrubbing. No differences in individual mean frame-wise displacement between the ICA (median 0.126 mm) and Control (median 0.114 mm) groups were demonstrated based on a Mann-Whitney U test (U = 536, p = .76), and no group differences in individual number of time-points with frame-wise displacement > 0.5 mm were demonstrated (ICA median = 2, Control median = 2; U = 515.5, p = .56), a threshold above which notable correlation changes have been observed (Power et al. 2014). Denoising of the functional data included removal of 5 principal components from white matter and five principal components from cerebrospinal fluid (Behzadi et al. 2007), linear detrending, bandpass filtering (0.01-0.1 Hz), and regression of the 6 realignment parameters and their first derivative, as well as regression of the volumes with a frame-wise displacement > 0.5 mm. All denoising steps were implemented in the CONN functional connectivity toolbox release 18b (Whitfield-Gabrieli and Nieto-Castanon 2012) and applied to both preprocessed spatially smoothed and preprocessed non-smooth data, as both were used in subsequent analyses.

#### Functional connectivity analyses

To assess potential effects of ICA on the intrinsic functional connectivity of the olfactory cortex, two sets of analyses were conducted. First, the functional connectivity within the olfactory network, as defined by core olfactory processing regions (Seubert et al. 2013), was assessed and compared between groups. Thereafter, a more specific analysis of primary olfactory cortex, defined as the regions receiving direct input from the olfactory bulb (Zhou et al. 2019), was conducted, in which homotopic connectivity and regional homogeneity was compared between groups. Analyses were conducted in MATLAB 2019b.

##### Functional connectivity between core olfactory regions

The olfactory network used here was defined as six regions of interest (ROI) thought to play a key role in olfactory processing. All regions were chosen based on a previously published activation likelihood estimation analysis (ALE, neuroimaging meta-analysis) of olfactory processing (Seubert et al. 2013): the bilateral piriform cortex, the bilateral orbitofrontal cortex, and the bilateral anterior insula. Based on the peak MNI-coordinates of these regions in the ALE-analysis (for the anterior insula, the mean of three peaks in each hemisphere), spherical ROIs with a 9 mm radius were created (Figure 1A); a method mimicking the ROI definition in previous publications on olfactory resting-state networks (Tobia et al. 2016; Lu et al. 2019). For each individual, mean time series from these six ROIs were extracted from denoised unsmoothed data; “smoothing” is achieved by averaging the signal within the ROI. Functional connectivity between the ROIs was computed using pairwise Pearson’s correlation and the correlation coefficients were thereafter Fisher’s z-transformed for statistical comparisons. The connectivity between each pair of ROIs was compared between the ICA and Control groups using an ANCOVA with age, sex, and scanner site included as nuisance covariates.

**Figure 1.**
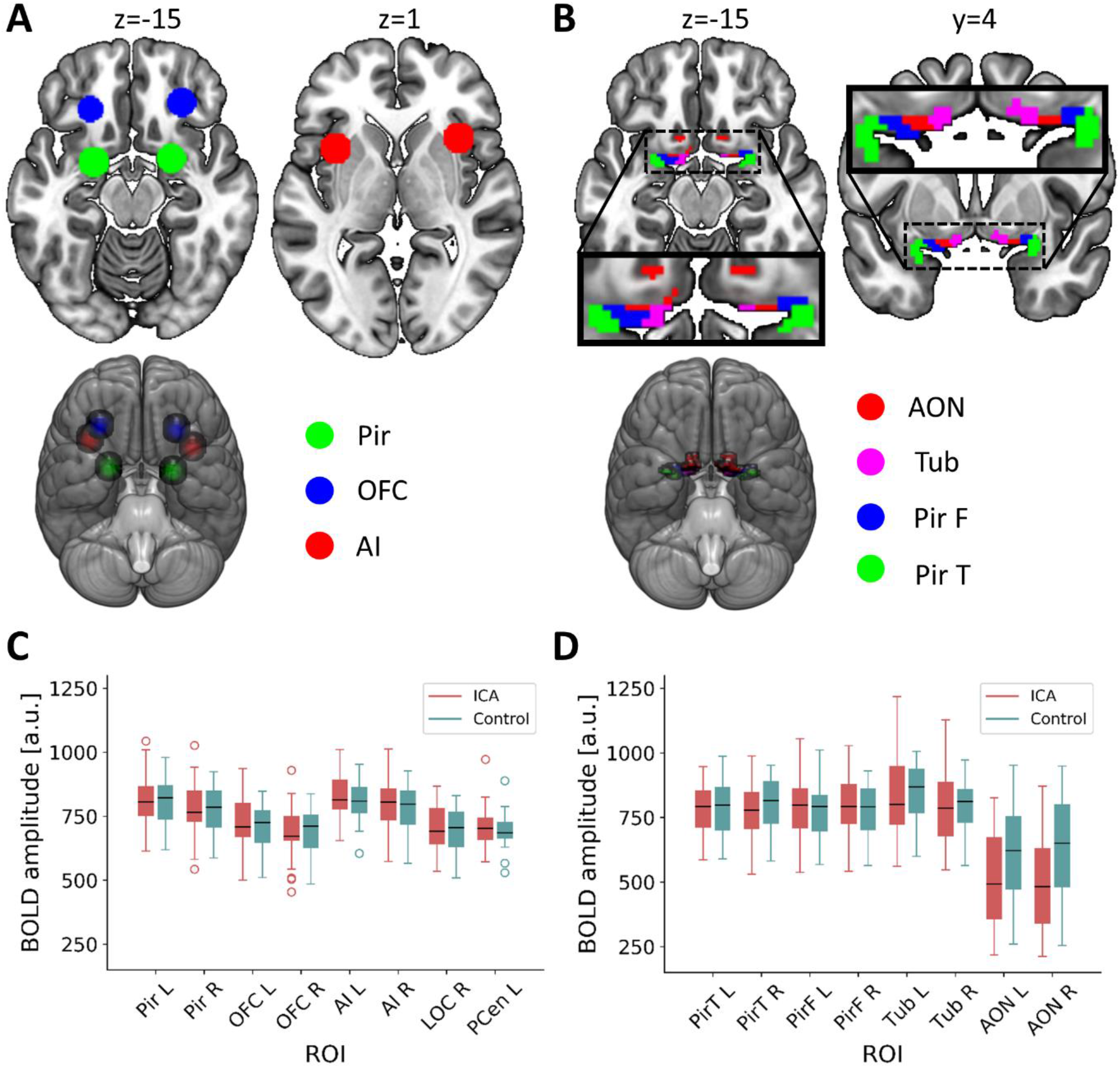
Definition of Regions of Interest (ROI) and their mean BOLD signal amplitudes. **A)** Core olfactory processing regions: spherical ROIs in bilateral piriform cortex (Pir; center coordinates [−22 0 −14] [22 2 −12]), orbitofrontal cortex (OFC; center coordinates [−24 30 −10] [28 34 −12]), and anterior insula (AI; center coordinates [−35 11 1] [35 17 1]). **B)** Primary olfactory subregions that together comprise the primary olfactory ROI (Zhou et al. 2019): anterior olfactory nucleus (AON), olfactory tubercle (TUB), frontal piriform cortex (PirF), and temporal piriform cortex (PirT). **C)** Boxplot of mean amplitude of preprocessed, but not denoised, BOLD signals in the six core olfactory processing ROIs from **A** and two non-olfactory reference ROIs where only small susceptibility artifacts is to be expected: left postcentral gyrus (PCen L; center coordinates [−44 −27 52]; Supplementary Figure S1A) and right lateral occipital cortex (LOC R; center coordinates [45 −74 4]; Supplementary Figure S1A). The borders of the boxes indicate the 1^st^ and 3^rd^ quartile, the whiskers stretch to the furthest data points within 1.5 interquartile range above/below the boxes, the black line indicates the 2^nd^ quartile (median); a.u. = arbitrary units. **D)** Boxplot of mean amplitude of preprocessed, but not denoised, BOLD signals in the eight primary olfactory subregions from **B**. A marked decrease in signal strength in the AON is visible. The borders of the boxes indicate the 1^st^ and 3^rd^ quartile, the whiskers stretch to the furthest data points within 1.5 interquartile range above/below the boxes, the black line indicates the 2^nd^ quartile (median); a.u. = arbitrary units.

##### Functional connectivity within primary olfactory cortex

We here use the primary olfactory cortex ROI according to the definition by Zhou and colleagues (2019), which includes eight subregions: the bilateral anterior olfactory nucleus, the bilateral olfactory tubercle, the bilateral frontal piriform cortex, and the bilateral temporal piriform cortex (Figure 1B). To assess potential functional deviations within primary olfactory cortex related to ICA, two different functional connectivity measures were used. First, local similarity of the BOLD signal was measured using regional homogeneity (Zang et al. 2004). Second, homotopic similarity of the BOLD signal, i.e., the correlation of the BOLD signal of corresponding regions in opposite hemispheres, was measured using voxel-mirrored homotopic connectivity (Zuo et al. 2010). Both connectivity measures were computed within the complete primary olfactory ROI, comprised of the eight subregions, using functions from the DPARSF package (Chao-Gan and Yu-Feng 2010). To avoid artificially increasing the similarity of BOLD signal in neighboring voxels, regional homogeneity was computed on non-smooth data with Kendall’s coefficient of concordance using 27 voxels (one center voxel and its 26 nearest neighbors). The resulting homogeneity maps were thereafter smoothed with an isotropic 6 mm full width at half maximum (FWHM) Gaussian smoothing kernel. For the voxel-mirrored homotopic connectivity analysis, the functional data was smoothed with an isotropic 6 mm FWHM Gaussian smoothing kernel prior to analysis. Correlation between each pair of sagittally mirrored voxels were computed and converted to Fisher’s z for statistical comparisons. Voxel-wise group comparison of regional homogeneity and voxel-mirrored homotopic connectivity were implemented in SPM12 using a GLM with age, sex, and scanner site as nuisance covariates. A statistical threshold of p < .05 family-wise error (FWE) corrected within the ROI was used for all analyses; if no voxel reached significance, a threshold of p < .01 was implemented to assess potential differences at a more liberal statistical threshold.

#### Data quality control

Two different measures were used to assess the quality of our data. First and foremost, the amplitude and variability of BOLD-signal in our ROIs were assessed given that the orbitofrontal cortex lies in close proximity to the sinuses and is therefore commonly affected by macroscopic susceptibility artifacts (Ojemann et al. 1997). In addition to our ROIs, two control regions in which we did not expect a presence of extensive signal distortion were included for comparison: 9 mm radius spheres in right lateral occipital cortex (LOC) and in left postcentral gyrus (Supplementary Figure S1A). For each individual, the mean time series from all 16 regions (six core olfactory processing regions, eight primary olfactory subregions, and two control regions) were extracted from preprocessed functional data prior to denoising (except for the removal of a linear slope). The BOLD time series were visually inspected and mean and standard deviation were computed for each individual and region. Signal strength was comparable for all regions except the bilateral AON, which showed a decreased amplitude, specifically noticeable in the ICA group but also clearly visible in the Control group (Figure 1C, 1D; Supplementary Figure S1B, S1C).

As a second quality control measure, in our control group, we set out to replicate the intrinsic olfactory network reported by Tobia and colleagues (2016), who based their network on a seed-to-voxel analysis with spherical seeds using the same orbitofrontal and piriform coordinates as were used by us. To strictly adhere to their method, in this analyses, we used an 8 mm3 smoothing kernel and conducted a seed-to-voxel analysis. Based on visual comparison of the resulting connectivity maps with those presented by Tobia and colleagues, and inclusion of the regions mentioned in their paper, such as thalamus and hippocampus, our results match those previously presented (Supplementary Figure S2, S3).

## RESULTS

### Preserved functional connectivity between olfactory regions in individuals with congenital anosmia

To investigate the effects of ICA on the functional connectivity between the defined core areas in olfactory processing, pairwise correlations between the six ROIs (bilateral orbitofrontal cortex, piriform cortex, and anterior insula) were computed for each individual and compared between groups (Figure 2). Within both the ICA and Control groups, high functional connectivity between corresponding regions in opposite hemispheres was found, especially for anterior insula and piriform cortex. However, we could not demonstrate significant differences between the ICA and Control group in connectivity between any of the ROIs (all p ≥ .306; for detailed statistical results, see Supplementary Table S1). A lack of statistical support for group differences is not, however, the same as statistical support for a lack of group differences.

**Figure 2.**
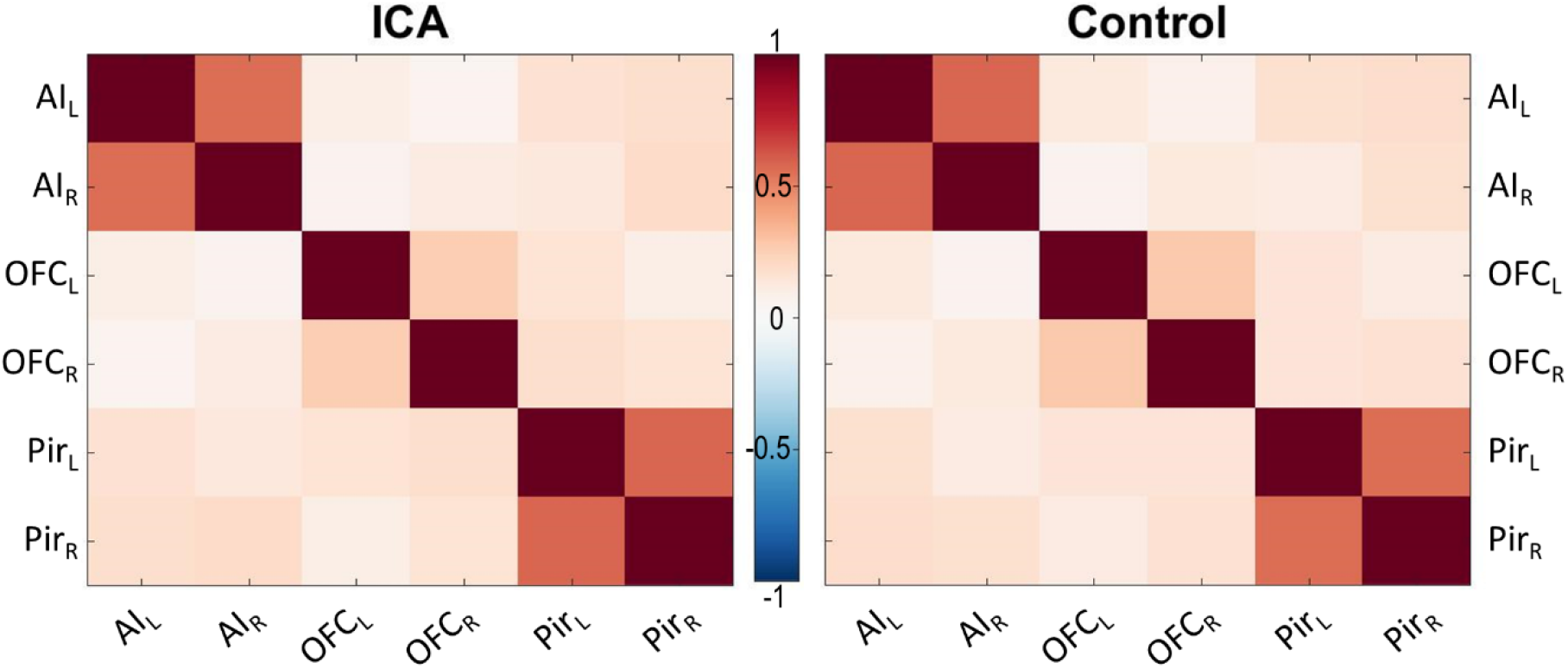
Correlation matrices for the olfactory network. No significant group differences in connectivity within the olfactory network outlined in Figure 1A at a p<.05, uncorrected, statistical threshold. AI = anterior insula, OFC = orbitofrontal cortex, Pir = piriform cortex; L=left and R=right hemisphere. Color bar denotes r-values.

To better assess whether these results could be interpreted in favor of the null hypothesis, i.e., no difference in connectivity between the two groups, Bayesian independent samples t-tests were conducted for the 15 connections using JASP (Version 0.11.1; https://jasp-stats-org). All tests yielded anecdotal to moderate evidence for the null hypothesis (2.64 < BF01 < 3.99; full results in Supplementary Table S2 and https://osf.io/3qsca/

### Lifelong absence of olfactory input does not alter regional homogeneity or homotopic connectivity in primary olfactory areas

To assess ICA-related functional deviations within the primary olfactory cortex, regional homogeneity and voxel-mirrored homotopic connectivity was compared between ICA and Control groups. Neither regional homogeneity (Figure 3A), nor voxel-mirrored homotopic connectivity (Figure 3C), was significantly different between groups at our pre-defined threshold of p < .05, FWE corrected within the ROI (Figure 3B). At the more liberal statistical threshold of p < .01, there were still no significant group differences in either one of the two connectivity measures. Because no significant group differences in either regional homogeneity or voxel-mirrored homotopic connectivity were demonstrated, further group comparisons using Bayesian independent samples t-tests were done, analogous with the analysis of the connectivity between the core olfactory regions. Specifically, for each individual, the mean homogeneity and homotopic connectivity were extracted from each of the eight primary olfactory subregions and compared between groups. All comparisons yielded anecdotal to moderate support for the null hypothesis (regional homogeneity: 2.02 < BF01 < 4; voxel-mirrored homotopic connectivity: 1.85 < BF01 < 3.99; Supplementary Table S3 and https://osf.io/3qsca/).

**Figure 3.**
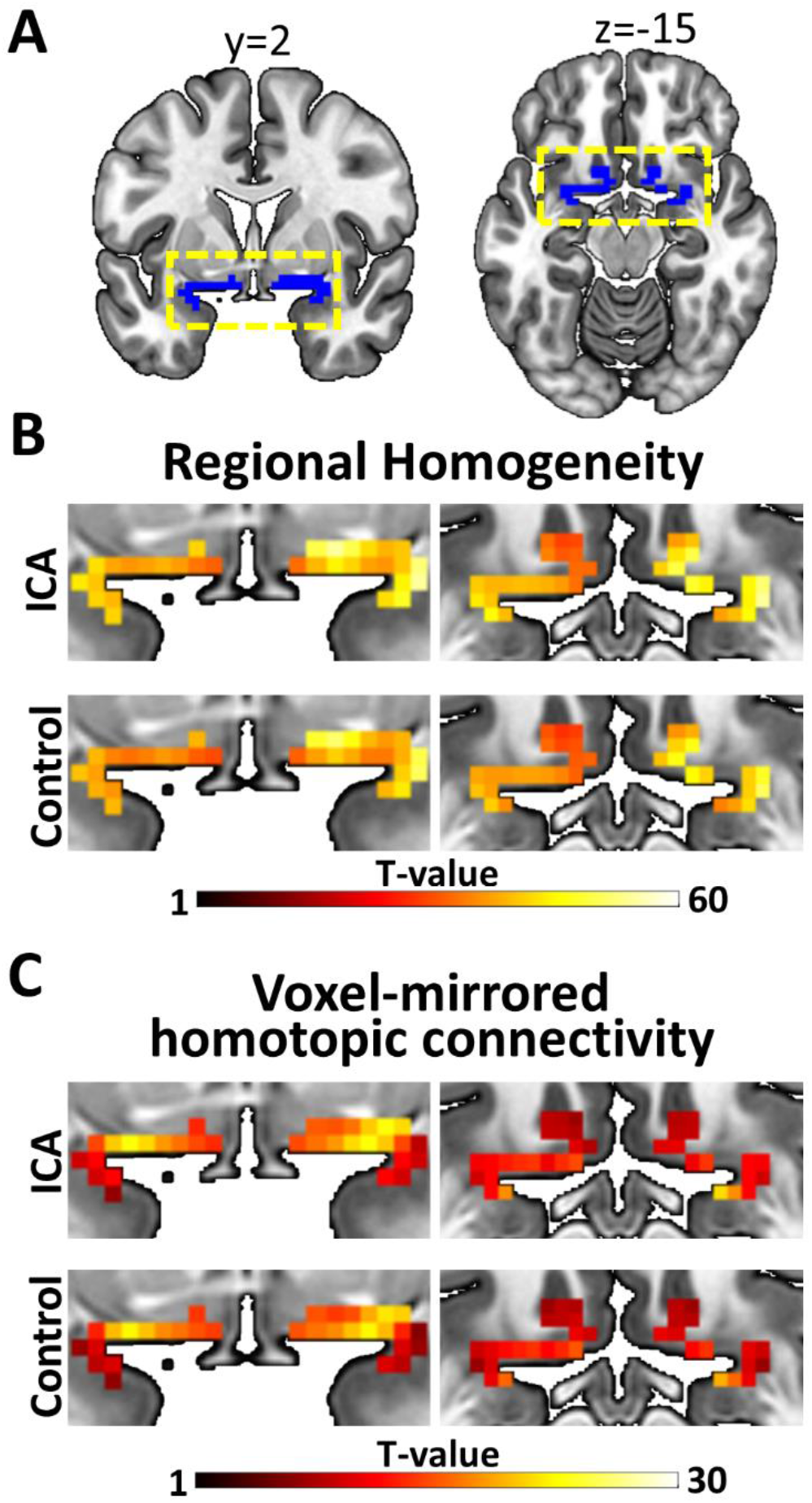
Connectivity results for the olfactory cortex. **A**) Primary olfactory ROI, for which the analyses were done, are marked in blue. The yellow dashed boxes mark the part of the cortex that is magnified in B and C. **B**) Regional homogeneity in both groups (color bar indicate t-values). No significant group differences at p<.01, uncorrected. **C**) Voxel-mirrored homotopic connectivity in both groups (color bar indicate t-values). No significant group differences at p<.01, uncorrected.

### Lifelong absence of olfactory input does not alter regional homogeneity or homotopic connectivity in primary olfactory areas

To assess ICA-related functional deviations within the primary olfactory cortex, regional homogeneity and voxel-mirrored homotopic connectivity was compared between ICA and Control groups. Neither regional homogeneity (Figure 3A), nor voxel-mirrored homotopic connectivity (Figure 3C), was significantly different between groups at our pre-defined threshold of p < .05, FWE corrected within the ROI (Figure 3B). At the more liberal statistical threshold of p < .01, there were still no significant group differences in either one of the two connectivity measures. Because no significant group differences in either regional homogeneity or voxel-mirrored homotopic connectivity were demonstrated, further group comparisons using Bayesian independent samples t-tests were done, analogous with the analysis of the connectivity between the core olfactory regions. Specifically, for each individual, the mean homogeneity and homotopic connectivity were extracted from each of the eight primary olfactory subregions and compared between groups. All comparisons yielded anecdotal to moderate support for the null hypothesis (regional homogeneity: 2.02 < BF01 < 4; voxel-mirrored homotopic connectivity: 1.85 < BF01 < 3.99; Supplementary Table S3 and https://osf.io/3qsca/).

## DISCUSSION

We investigated whether a lifelong absence of olfactory input is linked to altered function of core olfactory processing regions by comparing resting-state functional connectivity between individuals with isolated congenital anosmia (ICA) and matched controls. Specifically, the functional connectivity between core olfactory regions as well as the regional homogeneity and homotopic connectivity within primary olfactory cortex were assessed. In stark contrast to our hypotheses, none of our results support the notion that ICA is linked to atypical functional connectivity in the olfactory cortex. This implies that, unlike visual and auditory sensory deprivation, a life-long absence of olfactory sensations has limited effect on the functional connectivity of the core processing regions of the deprived sensory modality.

We found no difference between individuals with ICA and match healthy individuals in resting-state functional connectivity within the pre-defined olfactory network. Albeit a decrease in functional connectivity for ICA individuals was hypothesized, we suggest that the demonstrated lack of group differences are an indication of group similarity rather than an indication of inability to find existing differences due to, e.g., low power. We argue this based on the additional Bayesian analyses that showed support for the null hypotheses (no group differences). The fact that individuals with ICA, that have had a lifelong absence of olfactory input, did not demonstrate atypical connectivity raises interesting questions about the development of olfactory connectivity and, specifically, the influence of early olfactory input on the olfactory processing network.

Results from developmental studies (Zuo et al. 2010; Anderson et al. 2014) and studies investigating functional connectivity in visual sensory deprivation (A. Jiang et al. 2015; Liu et al. 2011; Hou et al. 2017; Watkins et al. 2012) suggest that higher homogeneity and lower homotopic connectivity would be expected in sensory processing regions lacking sensory input from birth. In contrast to these expectations, our results indicate that neither the regional homogeneity, nor the voxel-mirrored homotopic connectivity, in primary olfactory cortex is affected by the lack of olfactory input in individuals with ICA. Interestingly, the very same ICA individuals also demonstrated an unexpected lack of morphological reorganization in primary olfactory cortex (Peter et al. 2019), constituting an additional difference to blind individuals who demonstrate clear morphological alterations in primary visual cortex (Park et al. 2009; Ptito et al. 2008). However, because cortical morphology and regional homogeneity is related (L. Jiang et al. 2015; A. Jiang et al. 2015), the lack of altered functional connectivity within the primary olfactory cortex in individuals with ICA could logically follow upon the lack of morphological alterations in said region. Independent of the correspondence between the functional and structural results, the discrepancy between the effects of olfactory and visual sensory deprivation on sensory processing regions remains. Why a lifelong lack of olfactory input has little implications for the connectivity (and morphology) of primary olfactory regions, at least during rest, is yet to be determined.

In rodent models, experimental olfactory bulb ablation has different effects on the development of the piriform cortex depending on at what age the ablation occurs. Removal of the olfactory bulb right after birth, thus removing afferent input to the piriform (primary olfactory) cortex and mimicking congenital or very early olfactory sensory deprivation, has little to no effect on the cortical thickness of the piriform cortex, whereas a later removal leads to a definitive thinning of the piriform cortex (Friedman and Price 1986a; Friedman and Price 1986b; Westrum and Bakay 1986). The cortical thickness within the piriform cortex seems to be preserved in animals with early bulb removal because intracortical association fibers extend into the outer cortical layer normally occupied by the afferents from the olfactory bulb. Interestingly, there does not seem to be any extension of these association fibers into deafferented regions that they normally do not occupy (Friedman and Price 1986b). This finding indicates that, apart from the obvious lack of afferents from the olfactory bulb, structural connectivity remains virtually stable in very early olfactory sensory deprivation. This stability of structural connectivity in primary olfactory cortex demonstrated in non-human animal models could help explaining the absence of significant group differences in functional connectivity in the present study. In addition, the plasticity within piriform cortex in healthy rodents has been suggested to be more strongly regulated by input from the orbitofrontal (secondary olfactory) cortex than from the olfactory bulb (Strauch and Manahan-Vaughan 2017). Because the orbitofrontal cortex is a multimodal region, it could potentially provide a similar amount of input to piriform cortex independent of whether input from an olfactory bulb occurs or not, and thereby stimulate similar development of connectivity. The fact that piriform cortex has been associated with complex processes, such as attention and memory (Zelano et al. 2005; Zelano et al. 2009), and even processing of non-olfactory information such as visual (Porada et al. 2019) and intranasal trigeminal sensations (Albrecht et al. 2010), further supports the notion that piriform cortex might receive a comparable magnitude of neural input from neocortical areas even in the absence of olfactory input. One of the most noteworthy examples is the fact that piriform cortex is activated by odorless sniffs (Sobel et al. 1998), an activation that, at least tentatively, also occurs in individuals with congenital anosmia (Weiss et al. 2016). Based on our results combined with these structural and functional observations, we can speculate that olfactory input might not be crucial to develop and maintain normal connectivity within and between core olfactory regions when complete olfactory sensory deprivation is congenital or occurs very early in life.

Resting-state fMRI has led to a greater understanding of the functional organization of the human brain (Biswal et al. 1995; Fransson 2005); however, the method has limitations. First, data quality and analysis results are sensitive to subject motion during data acquisition (Power et al. 2012; Van Dijk et al. 2012). To decrease the effects of motion-related noise in our data, we used a number of preprocessing steps including, e.g., scrubbing of volumes with a frame-wise displacement exceeding 0.5 mm. Importantly, no significant differences were observed between groups in median frame-wise displacement or number of scrubbed volumes. This indicates that a potential difference in connectivity between groups is unlikely to be concealed by a potential group difference in motion. A second problem with BOLD fMRI is susceptibility artifacts that are particularly troublesome in orbitofrontal regions due to their close proximity to the sinuses (Ojemann et al. 1997). To estimate the effects of this potential signal loss problem, we assessed the amplitude of the raw BOLD signal in our olfactory ROIs and in two control ROIs where we would expect a lower degree of susceptibility artifacts. The signal amplitudes in our olfactory ROIs were comparable to the control regions’ with the exception of the bilateral AON (a subregion of our primary olfactory ROI), which showed noticeably lower amplitude in both groups. Hence, results from this particular primary olfactory subregion should be interpreted with caution. Based on both subject motion and BOLD signal strength, we argue that the signal quality is good in the present data. In addition, the fact that we were able to replicate the olfactory functional connectivity previously demonstrated by Tobia and colleagues (Tobia et al. 2016) further supports this notion and we maintain that the lack of group differences demonstrated here are unlikely to be caused by low data quality.

Although none of the group comparisons of functional connectivity indicated any differences between individuals with ICA and controls, it seems unlikely that a lifelong lack of olfactory input has absolutely no effect on the connectivity of core olfactory processing regions, even during rest. Needless to say, we are inherently limited by the spatial and temporal resolution of the methods we use and are thereby unable to draw conclusions regarding potential alterations in connectivity occurring at spatial or temporal scales not investigated here. Although we do not claim that a lifelong lack of olfactory input has absolutely no effect on the connectivity of the olfactory cortex, our results do imply that even if atypical connectivity (between or within olfactory regions) not detectable here exists in individuals with ICA, these potential alterations are far from the magnitude of those demonstrated in visual cortex of blind individuals where connectivity alterations are clearly detectable using ROI-to-ROI functional connectivity (Bauer et al. 2017), regional homogeneity (A. Jiang et al. 2015; Liu et al. 2011), and homotopic connectivity (Watkins et al. 2012; Hou et al. 2017).

The lack of group differences in resting-state functional connectivity between the core olfactory processing regions could be interpreted as a preserved functional olfactory network displayed by individuals with ICA despite a lifelong absence of olfactory input. However, although these core olfactory processing regions were chosen based on their known involvement in olfactory processing (Seubert et al. 2013) and previous use as seeds for an olfactory resting-state network (Tobia et al. 2016; Karunanayaka et al. 2017; Lu et al. 2019), it is important to note that even in our control group, consisting of individuals with an established normal sense of smell, the functional connectivity between these regions was not particularly high; high correlations were only demonstrated between bilateral regions within the network. Hence, the interpretation of our results as a preserved functional olfactory resting-state network in individuals with ICA seems less plausible, as our results challenge the idea of an olfactory resting-state network consisting of the proposed regions. Additionally, altered functional connectivity in individuals with acquired anosmia has been demonstrated during active sniffing, but not during normal breathing (Kollndorfer et al. 2015), and both auditory and visual sensory networks are included among the established resting-state networks (Damoiseaux et al. 2006; Power et al. 2011), present already in infants (Fransson et al. 2007), whereas an olfactory network is absent in these reports. This indicates that a clear olfactory network might not be detectable by BOLD fMRI during rest. Further investigation of olfactory connectivity during rest is warranted to more thoroughly establish the olfactory network in health.

To enable the inclusion of a greater number of individuals with the rare condition ICA, data was collected at two different locations. The use of multiple sites has been reported to have minimal effects on the reliability of functional connectivity, but uncontrolled differences across sites may potentially introduce bias (Noble et al. 2019). Thus, efforts were made to minimize the effects of scanning site: 3T Siemens scanners with identical scanning sequence protocols were used at both sites, site was included as a nuisance covariate in the analyses, and importantly, the matched control was always scanned at the same site as the individual with ICA, thus preventing inter-group effects of scanning site. Albeit these efforts might not completely remove the intra-group variability due to scanning site, we do believe that the benefits of the substantially increased sample size clearly outweigh the costs.

In summary, our data strongly suggests that individuals with ICA demonstrate typical resting-state functional connectivity between core olfactory processing regions as well as within primary olfactory cortex despite a lifelong lack of olfactory experience. This suggests that, in sharp contrast to the visual cortex, the olfactory cortex does not require early sensory input to develop and maintain functional connectivity. These results call for further studies on the basis of this suggested developmental difference between sensory processing regions in general, and the development of olfactory cortical regions in particular.

## CONFLICT OF INTEREST

The authors have no conflict of interest to report.

## ACKNOWLEDGEMENTS

This work was supported by the Knut and Alice Wallenberg Foundation (KAW 2018.0152 to J.N.L.) and the Swedish Research Council (2017-02325 to J.N.L.).

## Notes

https://osf.io/3qsca/

### Competing Interest Statement

The authors have declared no competing interest.

