## Supplementary for "Normal Olfactory Functional Connectivity Despite Life-Long Absence of Olfactory Experiences"

Peter, M.G. *et al.* 2020

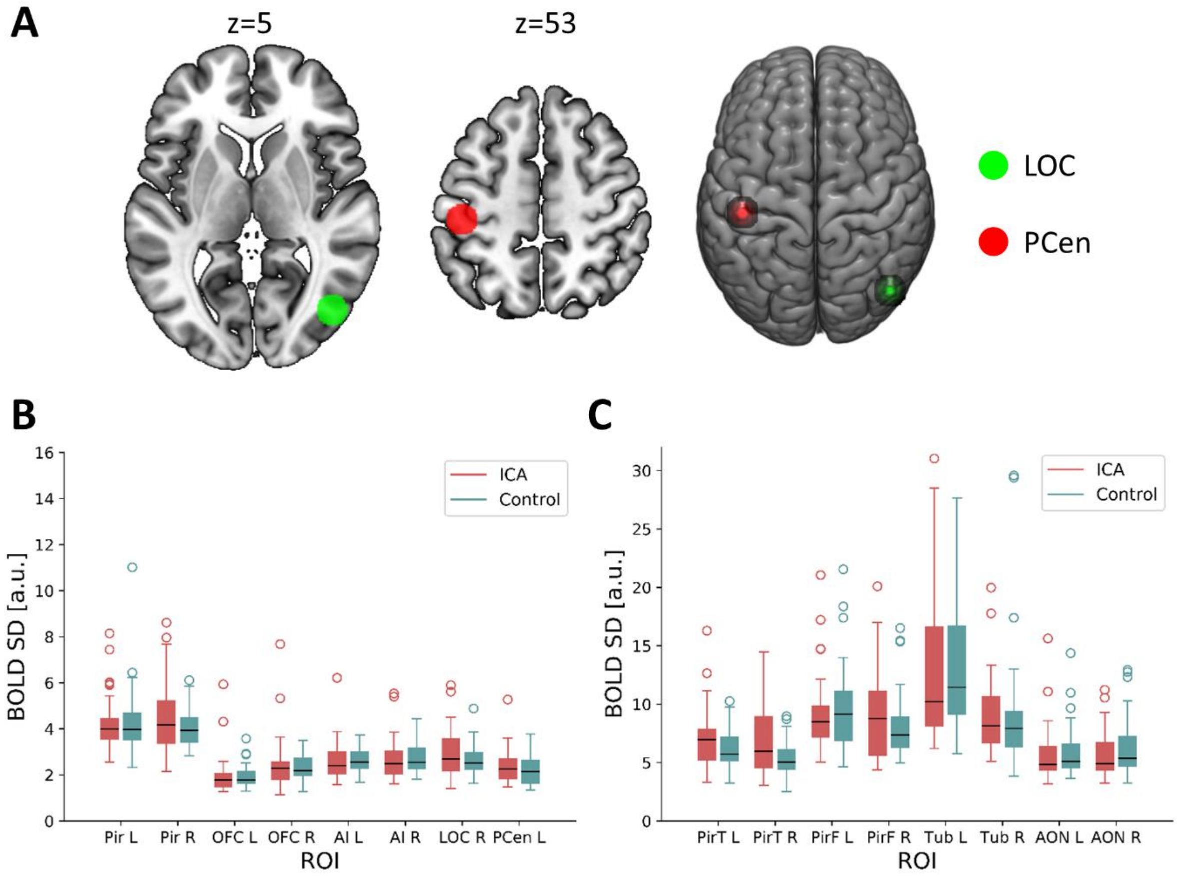


***Supplementary Figure S1 Reference regions and standard deviation of the BOLD time series in the regions of interest. A)*** *Two reference ROIs: LOC=lateral occipital cortex, PCen=postcentral gyrus**.** **B)** Boxplots of the standard deviations of the preprocessed, but not denoised, BOLD timeseries in core olfactory processing regions: piriform cortex (Pir), orbitofrontal cortex (OFC), and anterior insula (AI) **C)** Boxplots of the standard deviations of the preprocessed, but not denoised, BOLD timeseries in primary olfactory subregions: anterior olfactory nucleus (AON), olfactory tubercle (TUB), frontal piriform cortex (PirF), and temporal piriform cortex (PirT). Note that the primary olfactory subregions are much smaller than the spherical regions (about one tenth of the volume) and located close to the gray matter/cerebrospinal fluid border, which likely contributes to the larger standard deviations as an effects of noise and motion. The borders of the boxes indicate the 1st and 3rd quartile, the whiskers stretch to the furthest data points within 1.5 interquartile range above/below the boxes, the black line indicates the 2nd quartile (median); a.u. = arbitrary unit.*

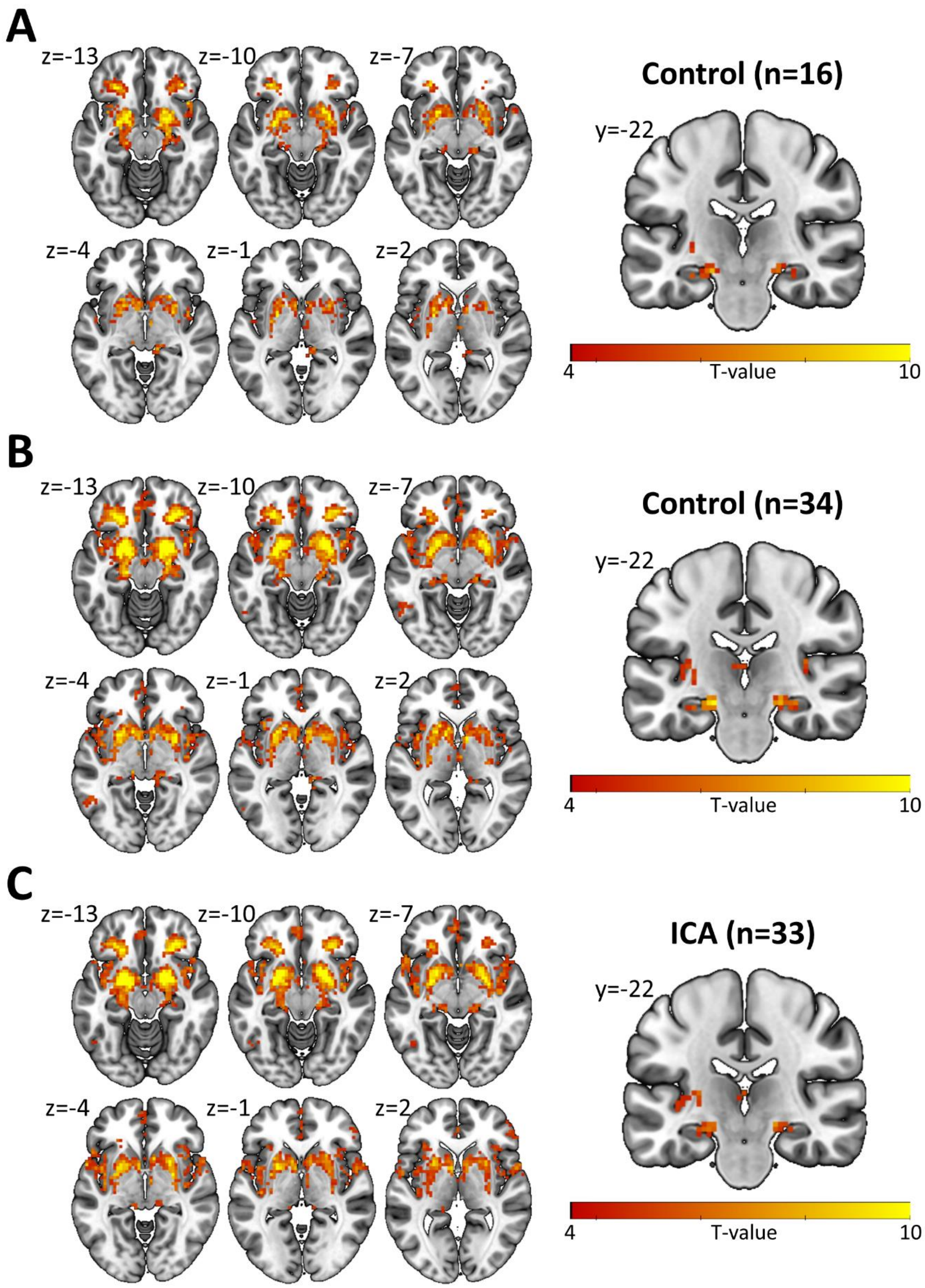


***Supplementary Figure S2 Resting-state functional connectivity from bilateral piriform and orbitofrontal seeds****. These connectivity maps are visually similar to the 'olfactory network' presented by Tobia and colleagues (1) , and include many of the same regions, e.g., the thalamus, medial prefrontal cortex, parahippocampal gyrus, and hippocampus.* ***A)*** *Connectivity in 16 control subjects to replicate the analysis by Tobia and colleagues (1) with the same sample size. Voxel-wise thresholded at p<.001 with a minimum cluster size of 20 voxels.* ***B)*** *Connectivity in Control group. Voxel-wise thresholded at a false discovery rate (FDR) of <.01 with a minimum cluster size of 20 voxels.* ***C)*** *Connectivity in ICA group. Voxel-wise thresholded at a false discovery rate (FDR) of <.01 with a minimum cluster size of 20 voxels.*

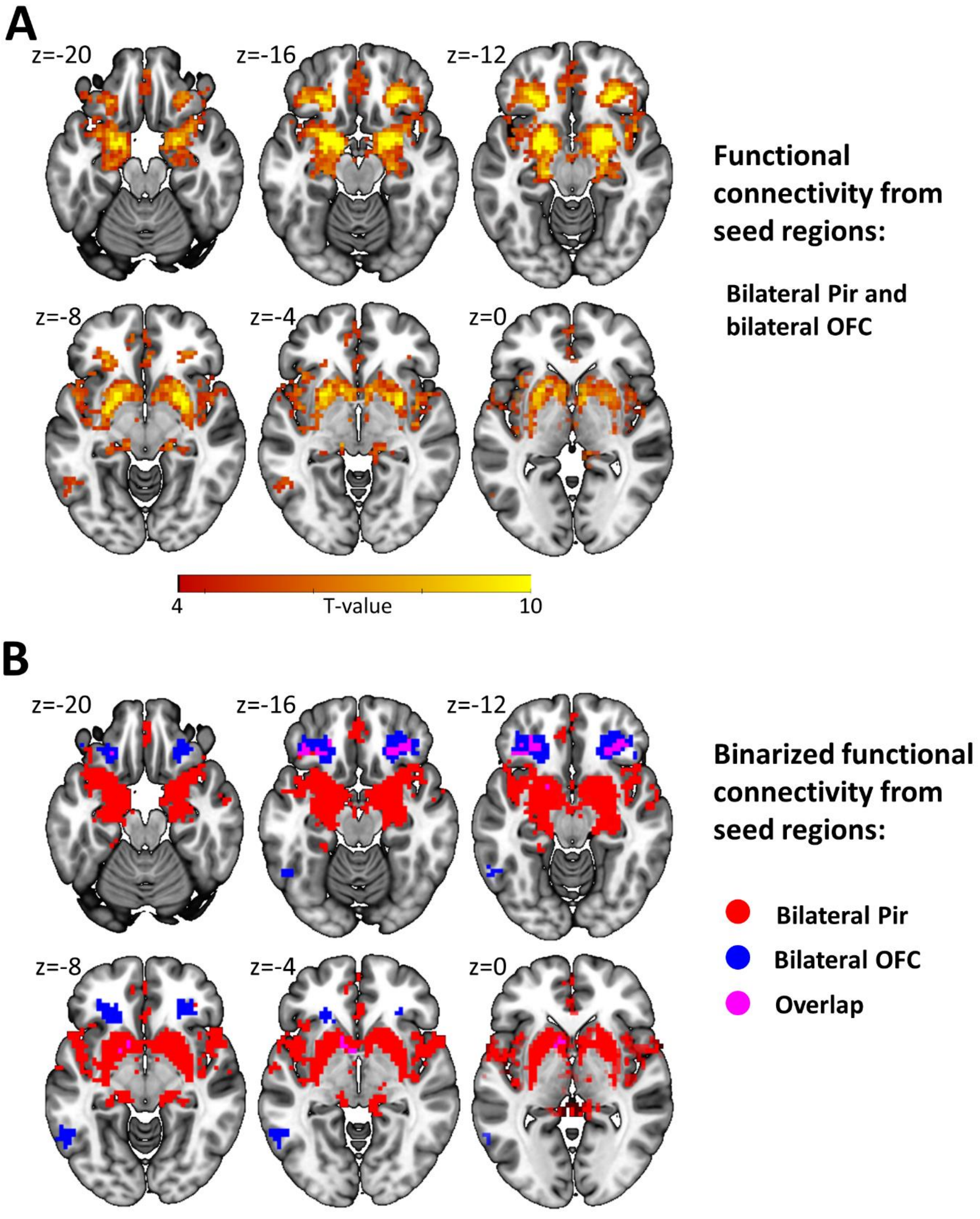


***Supplementary Figure S3. Resting-state functional connectivity from bilateral piriform and orbitofrontal seeds in the Control group*. *A)*** *The same connectivity map as presented in Supplementary Figure S1B (using bilateral orbitofrontal cortex and bilateral piriform cortex as seed regions; voxel-wise thresholded at FDR < .01 and minimum cluster size of 20 voxels), but displayed on different slices visualize the regions around the seeds (centers -14 ≤ z ≤ -10)* ***B)*** *The connectivity from bilateral piriform cortex (Pir; red color) and bilateral orbitofrontal cortex (OFC; blue color) visualized separately with overlap (violet color). Note that the overlap is not extensive, suggesting that the definition of this as one network should be further investigated.*

**Table S1.**

*Results for factor Group in ANCOVA of connectivity between core olfactory regions (age, sex, and scanner site as nuisance covariates)*

| Region 1 | Region 2 | *F(1,62)* | *p* |
|---|---|---|---|
| AI left | AI right | 0.37 | .544 |
| AI left | OFC left | 0.42 | .522 |
| AI left | OFC right | 0.16 | .69 |
| AI left | Pir left | 0.02 | .875 |
| AI left | Pir right | 0.05 | .819 |
| AI right | OFC left | 0.01 | .938 |
| AI right | OFC right | 0.01 | .923 |
| AI right | Pir left | 0.49 | .489 |
| AI right | Pir right | 1.07 | .306 |
| OFC left | OFC right | 0.57 | .454 |
| OFC left | Pir left | 0.1 | .748 |
| OFC left | Pir right | 0.36 | .551 |
| OFC right | Pir left | 0.79 | .378 |
| OFC right | Pir right | 0.06 | .805 |
| Pir left | Pir right | 0.43 | .512 |

AI = anterior insula, OFC = orbitofrontal cortex, Pir = piriform cortex

**Table S2.**

*Bayes factor for Bayesian independent samples t-test of functional connectivity between groups*

| Region 1 | Region 2 | $BF_{01}$ |
|---|---|---|
| AI left | AI right | 3.446 |
| AI left | OFC left | 3.325 |
| AI left | OFC right | 3.702 |
| AI left | Pir left | 3.940 |
| AI left | Pir right | 3.883 |
| AI right | OFC left | 3.980 |
| AI right | OFC right | 3.975 |
| AI right | Pir left | 3.267 |
| AI right | Pir right | 2.644 |
| OFC left | OFC right | 3.079 |
| OFC left | Pir left | 3.839 |
| OFC left | Pir right | 3.410 |
| OFC right | Pir left | 2.878 |
| OFC right | Pir right | 3.895 |
| Pir left | Pir right | 3.443 |

$BF_{01}$ = Bayes Factor in support of null hypothesis over alternative hypothesis, AI = anterior insula, OFC = orbitofrontal cortex, Pir = piriform cortex

**Table S3.**

*Bayes factor for Bayesian independent samples t-test of regional homogeneity and voxel-mirrored homotopic connectivity between groups*

| Region | ReHo $BF_{01}$ | VMHC $BF_{01}$ |
|---|---|---|
| AON left | 2.858 | 2.974 |
| AON right | 2.026 | 3.323 |
| Pir F left | 3.754 | 2.953 |
| Pir F right | 3.991 | 3.111 |
| Pir T left | 3.976 | 2.758 |
| Pir T right | 3.906 | 1.857 |
| Tub left | 3.717 | 3.438 |
| Tub right | 3.958 | 3.980 |
| Prim Olf | 3.525 | 3.646 |

ReHo = regional homogeneity, VMHC = voxel-mirrored homotopic connectivity, $BF_{01}$ = Bayes Factor in support of null hypothesis over alternative hypothesis, AON = anterior olfactory tubercle, Pir F = frontal piriform cortex, Pir T = temporal piriform cortex, Tub = olfactory tubercle, Prim Olf = all primary olfactory subregions together
